# Endoplasmic reticulum stress engenders immune-resistant, latent pancreatic cancer metastases

**DOI:** 10.1101/187484

**Authors:** Arnaud Pommier, Naishitha Anaparthy, Nicoletta Memos, Z Larkin Kelley, Alizée Gouronnec, Jean Albrengues, Mikala Egeblad, Christine A. Iacobuzio-Donahue, Scott K. Lyons, Douglas T. Fearon

## Abstract

Patients who have had their primary pancreatic ductal adenocarcinoma (PDA) surgically resected often develop metastatic disease, exemplifying the problem of latent metastases. Livers from patients and mice with PDA contained single, disseminated cancer cells (DCCs) with an unusual phenotype of being cytokeratin-19 (CK19)^-^ and MHC class I (MHCI)^-^. We created a mouse model to determine how DCCs develop, their relationship to metastatic latency, and the role of immunity. Intra-portal injection of immunogenic PDA cells into pre-immunized mice seeded livers only with single, non-replicating DCCs lacking MHCI and CK19; naïve recipients had macro-metastases. Transcriptomic analysis of PDA cells with the DCC phenotype demonstrated an endoplasmic reticulum (ER) stress response. Relieving ER stress with a chemical chaperone, in combination with T cell-depletion, stimulated outgrowth of macro-metastatic lesions containing PDA cells expressing MHCI and CK19. The ER stress response is the cell-autonomous reaction that enables DCCs to escape immunity and establish latent metastases.

**One sentence summary:** Latent pancreatic cancer metastases are created when T cells select disseminated cancer cells in which immune resistance and quiescence have been imposed by endoplasmic stress.

## Introduction

Pancreatic ductal adenocarcinoma (PDA) is the fourth most common cause of death from cancer in the United States, and has a five year survival rate of 6% (*1*). This dismal prognosis is due in part to diagnoses not being made until the disease has extended beyond the primary tumor site so that surgery with the intention of cure may be conducted in only the 20% of patients who have no clinical evidence of local invasion or distant metastasis. Unfortunately, approximately 75% of these patients develop metastatic disease within two years after resection of their primary tumors (*2*, *3*), despite intra-operative examination of the liver confirming the absence of macro-metastatic lesions in over 80% of surgical patients (*4*). These observations lead to the conclusion that latent metastases, detectable only microscopically, were present in these patients and were responsible for the post-operative development of metastatic disease.

Latent metastases having a potential for outgrowth had been considered to represent lesions in which cancer cell proliferation is balanced by immune-mediated cancer cell death (*5*-*7*), but a more recent explanation invokes quiescent, single disseminated cancer cells (DCCs) (*8*-*10*). Single, non-replicating DCCs have been observed in several cancer types, most often in the bone marrow (*11*, *12*), but whether quiescence is enforced by the microenvironment or is cancer cell-autonomous is not known (*13*). Immunity, both innate (*14*) and adaptive (*6*, *15*, *16*), also is likely to have a role in the selection and/or maintenance of latent DCCs, as has long been suspected based on donor-derived cancer occurring in immune suppressed recipients of allografts (*17*, *18*). However, there is an unexplained paradox of immunity preventing the outgrowth of latent metastases while not eliminating latent metastases.

In the present study, we examine the problem of latent metastases in PDA by developing a mouse model that replicates the characteristics of hepatic DCCs in human and spontaneous mouse PDA. We have studied the metastatic process in the context of an on-going adaptive immune response because of the occurrence of cancer cell-specific immunity in human and mouse PDA (*19*-*22*). We find that a PDA-specific adaptive immune response selects DCCs in which the ER stress response accounts for both quiescence and resistance to immune elimination. Accordingly, outgrowth of DCCs to macro-metastases requires not only relief from the cancer cell-autonomous ER stress response, but also suppression of systemic immunity.

## Results

### Quiescent, single disseminated cancer cells in the livers of human and mice with PDA

To determine whether hepatic DCCs occur in human PDA, we microscopically examined tissue sections from the primary tumors and livers of five patients with PDA who had no clinically detectable metastases. The tumors were genotyped as having p53 loss-of-heterozygosity, which permitted staining for mutant p53 accumulation as an identifier of cancer cells. p53^+^ cancer cells were present in both the primary tumors and livers of all five patients. The p53^+^ cancer cells resided in the livers as single cells that were consistently CK19^-^, Ki67^-^, and MHCI^-^, in contrast to the cancer cells in the primary tumors, which exhibited all three markers **(Fig. 1A-C)**. We also examined the livers from mice bearing the autochthonous LSL-Kras^G12D/+^; LSL-Trp53^R172H/+^; Pdx-1-Cre; Rosa^YFP^ (KPCY) model of PDA (*23*-*25*) that recapitulates human PDA. In livers devoid of macro-metastases, we found both YFP^+^ micro-metastases and DCCs. While the micro-metastases always expressed CK19, Ki67 and MHCI, the single DCCs were mainly CK19^-^ (32/40), Ki67^-^ (22/22), and MHCI^-^ (28/28) **(Fig.1D-F)**. Therefore, the livers of patients and mice with PDA contain DCCs that share an unusual phenotype linking the loss of epithelial gene expression and quiescence with a potential for escape from T cell recognition.

**Figure 1:**
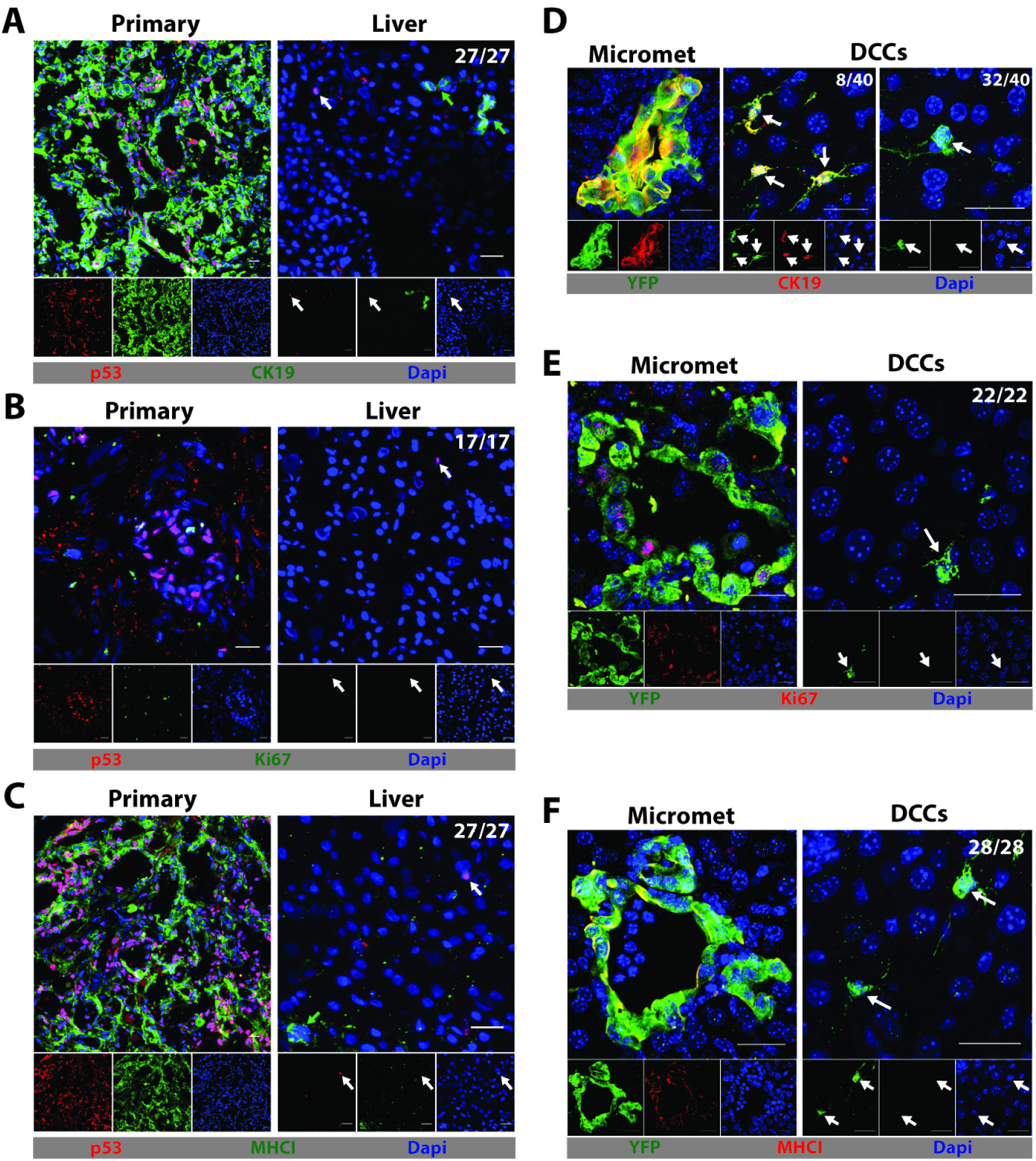
The presence of single DCCs having a characteristic phenotype in the livers of humans and mice with PDA. (A-C) IF of sections from the primary tumor and liver of a patient with PDA that have been stained with anti-p53 to reveal cancer cells (red) and (A) anti-CK19 (green), (B) anti-Ki67 (green) or (C) anti-MHCI (green). Photomicrographs are representative of five patients. (D-F) IF of sections from a liver of a KPCY mouse with spontaneous PDA and no hepatic macro-metastases. Sections were stained with anti-YFP to reveal cancer cells (green) and (D) anti-CK19 (red), (E) anti-Ki67 (red), and (F) anti-MHCI (red). Photomicrographs are representative of three mice. Frequency of events within the human and mouse DCC populations is indicated in the top right corner of the photomicrographs. White arrows designate DCCs and, in the sections of human livers, green arrows designate normal, liver-resident CK19^+^ or MHCI^+^ cells. Scale bar = 25μm.

### A model of hepatic metastasis in the context of an adaptive immune response

The absence of expression of MHCI on DCCs and the occurrence of cancer-specific CD8^+^ T cells in the genetically engineered mice model of PDA (*19*) suggested that DCCs may be selected by the immune response during the metastatic process. Accordingly, we developed a model of hepatic metastases that could assess the effect of a pre-existing immune response. A cell line was derived from a spontaneous liver metastasis of a mouse bearing an autochthonous PDA, and was stably transfected with a transposon vector directing the expression of diphtheria toxin receptor (DTR), *Herpes simplex* thymidine kinase (HSV-TK), firefly luciferase, and mTagBFP2 to generate mM1DTLB cells. Syngeneic C57Bl/6 mice were injected subcutaneously with 10^6^ mM1DTLB cells, tumors were grown for 14 days, and then eliminated by treatment with diphtheria toxin (DTx) and ganciclovir (GcV). These “pre-immunized” mice, and naïve mice were challenged by intra-splenic injections of 10^6^ mM1DTLB cells, followed by splenectomy, thereby seeding the liver *via* the portal vein, as in PDA **(Fig. 2A)**. In naïve mice, whole body bioluminescence increased from day 1 after injection, consistent with the growth of hepatic metastases. In pre-immunized mice, however, whole body bioluminescence decreased after day 1, and by day 7 luminescence was at background levels **(Fig. 2B, C)**. In additional cohorts of naïve and pre-immunized mice, livers were removed at intervals after the intra-splenic injection of mM1DTLB cells, and assessed for bioluminescence. Both photon flux **(Fig. 2B)** and visually detectable metastases increased in the livers of naïve mice between day 5 and 20, while metastatic foci were barely detectable in pre-immunized mice at day 5 and absent at later time points **(Fig. 2D)**. A potential role for T cells in the elimination of mM1DTLB cancer cells was suggested by the finding that in the livers of pre-immunized mice, tumor cells were frequently surrounded by both CD8^+^ and CD8^-^ CD3^+^ T cells by 24h **(Fig. S1A)**. This possibility was confirmed by treating pre-immunized mice with depleting antibodies to CD4 and CD8, alone or together, or with isotype control antibody **(Fig. S1B)**. Depleting either CD4^+^ or CD8^+^ T cells abrogated the effect of pre-immunization on suppressing the development of macro-metastases **(Fig. S1B)**, confirming this role of T cells.

**Figure 2:**
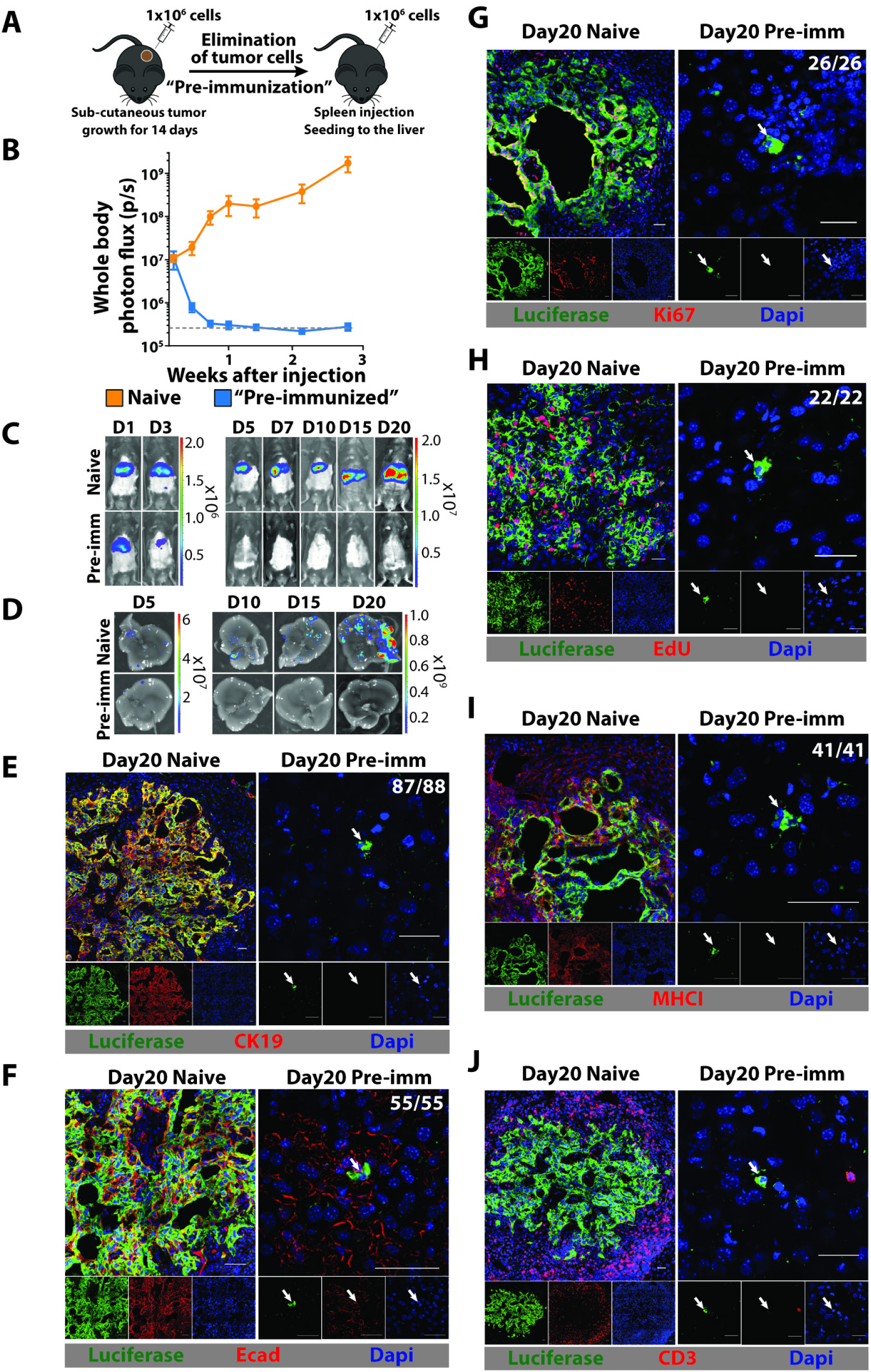
A mouse model for hepatic DCCs. (A) Mice are pre-immunized by subcutaneously injecting 10^6^ mM1DTLB PDA cells derived from a hepatic metastasis of a KPC mouse. After two weeks growth, tumors are eliminated by treating mice with DTx and GcV. For hepatic metastases, 10^6^ mM1DTLB PDA cells are injected intra-splenically into naïve and pre-immunized mice, followed immediately by splenectomy. (B, C) Tumor growth was measured by whole body bioluminescent imaging. (D) Ex vivo photon flux of whole livers was measured at day 5, 10, 15 and 20 after cancer cell injection. Results are representative of three experiments with at least five mice per group. Dashed gray line represents the background luminescence in tumor-free mice. (E-J) IF of sections from a liver of a naive mouse (left panels) or a pre-immunized mouse (right panels) that have been stained with anti-luciferase (green) to identify cancer cells and (E) anti-CK19 (red), (F) anti-Ecad (red), (G) anti-Ki67 (red), (H) EdU (red), (I) anti-MHCI (red), and (J) CD3 (red). For EdU staining, mice were injected every 12 hours with EdU for three days. Photographs are representative of 20 mice from three independent experiments. Frequency of events is indicated in the top right corner of the photomicrographs. White arrows designate DCCs. Scale bar = 25μm.

We also examined livers from naïve and pre-immunized mice microscopically, which revealed the presence of macro-metastatic lesions in the former, but only single DCCs in the latter. The DCCs differed from cancer cells in the macro-metastases by not expressing the epithelial markers, CK19 and E-cadherin (Ecad), not expressing Ki67 or incorporating EdU, and lacking MHCI **(Fig. 2E-I)**. Importantly, this phenotype of the DCCs of the metastasis model is similar to that of DCCs in human PDA and the KPCY mouse. T cells in the vicinity of DCCs were infrequent, whereas they surrounded macro-metastatic lesions in naïve mice **(Fig. 2J)**. The absence of CK19 and Ecad expression did not indicate an epithelial-mesenchymal transition (EMT) because DCCs did not express the EMT markers, desmin, αSMA, Snail1, or Slug **(Fig. S2)**. When naïve and pre-immunized mice were challenged with mM1DTLB cells *via* the tail vein, lung macro-metastases developed only in the naïve mice, whilst CK19^-^ DCCs were observed in the lungs of pre-immunized mice **(Fig. S3).** We also assessed the 1242 cell line derived from a primary PDA tumor of a KPC mouse, which had been modified with the same transposon vector, for its association with DCCs. When naïve and pre-immunized mice were challenged by intra-splenic injection of 1242DTLB cells, hepatic macro-metastases developed in the naïve mice, whilst only CK19^-^ DCCs were observed in the livers of pre-immunized mice **(Fig. S4).** To assess more stringently the proliferation of DCCs, we labelled the mM1DTLB cells with CFSE before intra-splenic injection into naïve or pre-immunized mice. Whereas all cancer cells in the macro-metastases of naïve mice had become CFSE^-^, DCCs in pre-immunized mice had retained CFSE **(Fig. S5A)**. Mice that had received intra-splenic injections of mM1DTLB cells were also given EdU in the drinking water for 20 days. While cancer cells in macro-metastases in naïve mice had incorporated EdU, almost all DCCs in pre-immunized mice were EdU^-^ **(Fig. S5B)**. Therefore, the occurrence of quiescent, MHCI^-^ DCCs in the absence of macro-metastases is a consequence of an on-going cancer-specific immune response.

### A latent capacity for outgrowth of DCCs is controlled by T cells

We examined whether a latent capacity of DCCs for outgrowth into macro-metastatic lesions might be revealed by T cell depletion. When depleting anti-CD4 and anti-CD8 antibodies were administered to pre-immunized mice three weeks after the establishment of DCCs, macro-metastases appeared in 10/15 mice. When T cells were depleted at nine weeks, macro-metastases appeared in only 2/15 mice **(Fig. 3)**. The metastases were composed of cancer cells that had re-expressed CK19 and MHCI, suggesting that DCCs must revert to an epithelial phenotype to initiate the formation of macro-metastases. The lower frequency of macro-metastases in mice in which T cells were depleted at nine weeks suggests either that DCCs with a capacity for reversion had decreased between three and nine weeks, or that individual DCCs lose this function with the passage of time. In favor of the former possibility is the finding that livers of pre-immunized mice have fewer DCCs at nine weeks than at three weeks **(Fig. S6A)**. This loss of DCCs may reflect the killing by T cells of spontaneous revertants, as suggested by the occasional occurrence of a CK19^+^ DCC surrounded by T cells in the pre-immunized mice **(Fig. S6B)**. The re-expression of MHCI by these growing metastases also suggests the means by which T cells control the outgrowth of spontaneously reverting DCCs. To demonstrate that T cells alone are both necessary and sufficient for controlling the growth of MHCI-DCCs, we depleted NK cells by administering anti-NK1.1 at the time of mM1DTLB cell challenge. Pre-immunized mice lacking NK cells were indistinguishable, with respect to the occurrence of DCCs and absence of macro-metastases, from control antibody-treated mice **(Fig. S7)**. This evidence for a dominant role of the T cell in controlling DCCs is supported by the finding that DCCs were never seen to be in contact with CD45^+^ **(Fig. S8A)**, F4/80^+^ **(Fig. S8B)**, CD19^+^ **(Fig. S8C)**, CD31^+^ **(Fig. S8D)** or αSMA^+^ **(Fig. S8E)** cells, the distribution of these cell types being similar to that in the normal liver **(Fig. S8 F-L)**.

**Figure 3:**
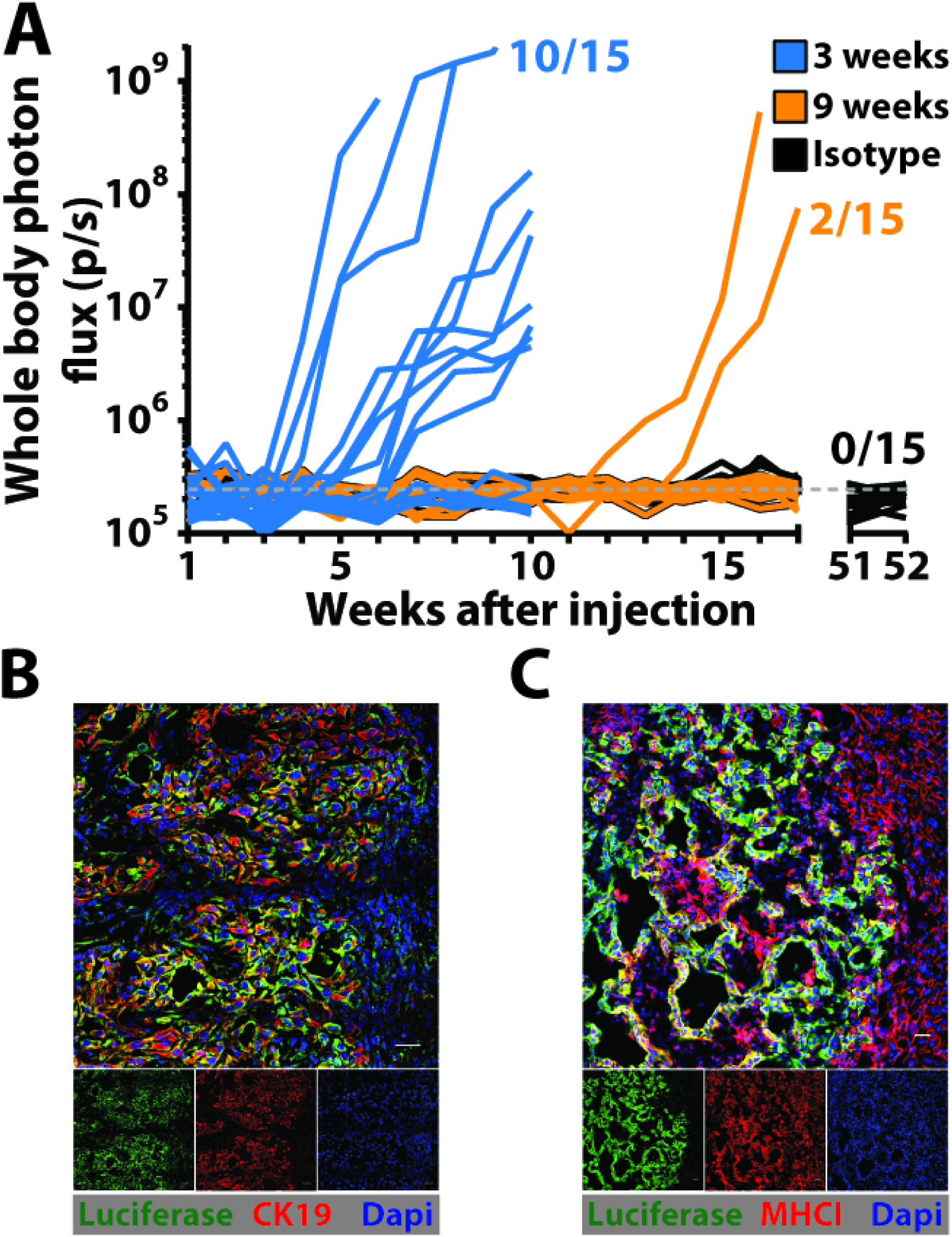
T cell control of outgrowth of latent DCCs. (A) The growth of hepatic metastases in pre-immunized mice that had been depleted of T cells by administration of antibodies to CD4 and CD8 beginning at three weeks or nine weeks after splenic injection of mM1DTLB PDA cells was assessed by bioluminescence imaging. One group of mice was also treated with isotype control antibody. (B and C) IF of sections containing macro-metastases from a liver of a pre-immunized mouse that had been depleted of T cells three weeks after splenic injection of mM1DTLB PDA cells. Anti-luciferase identifies cancer cells. Scale bar = 25μm.

### A rare sub-population of PDA cells in vitro with the phenotype of DCCs

The absence of MHCI expression by DCCs raised the possibility that these cells were present in the injected PDA population and were negatively selected by T cells. Indeed, approximately 1% of the mM1DTLB cells in tissue culture were Ecad^-^ and CK19^-^, and all Ecad^-^ cells were MHCI^-^ **(Fig. 4A, B)**. The Ecad^-^ cells resided in a non-proliferating sub-population of cells, as indicated by the resistance of these HSV-TK-expressing cells to GcV **(Fig. 4C)**. A phenotypic plasticity of the mM1DTLB cells was shown by culturing FACS-purified Ecad^+^ and Ecad^-^ cells for three days, and finding that they generated Ecad^-^ and Ecad^+^ cells, respectively **(Fig. 4D)**. The Ecad^-^ MHCI^-^ phenotype shared by PDA cells *in vitro* and the DCCs *in vivo* suggested that the former may be the precursors of the latter. We assessed this possibility by intra-splenically injecting 10^6^ Ecad^+^ and 10^4^ Ecad^-^ cells, respectively, into naïve and pre-immunized mice. The growth of Ecad^+^ macro-metastases in the livers of the naïve mice receiving Ecad^+^ cells was similar to that of naïve mice receiving unsorted mM1DTLB cells **(Fig. 4E)**. Microscopic examination of the livers of these mice, however, revealed no Ecad-DCCs **(Fig. 4F)**. Injecting Ecad^+^ cells in pre-immunized mice resulted in a rapid decline in photon flux, with no specific signal by day 7 **(Fig. 4E)**, and microscopic examination of these livers also demonstrated an absence DCCs **(Fig. 4F)**. Injecting Ecad^-^ cells into naïve mice resulted eventually in the growth of hepatic metastases **(Fig. 4E)**, and microscopy revealed the presence of DCCs **(Fig. 4F)**. Injection of Ecad^-^ cells into pre-immunized mice led to the occurrence only of DCCs **(Fig. 4E, F)**. The macro-metastases found in naïve mice injected with Ecad^-^ cells were CK19^+^ **(Fig. 4G)** and MHCI^+^ **(Fig. 4H)** indicating reversion to an epithelial phenotype. In summary, the origin of the DCC is the Ecad^-^ cell, as DCCs were not present in naïve or pre-immunized mice following the injection of Ecad^+^ cells. Reversion from the quiescent, Ecad^-^ state to the proliferating, Ecad^+^ phenotype is observed in naïve mice, but is masked by the on-going T cell response in pre-immunized mice because of the linkage of reversion to re-expression of MHCI. Therefore, the two states for mM1DTLB cells that are observed *in vivo*, the proliferating MHCI^+^/Ecad^+^/CK19^+^ macro-metastasis and the quiescent MHCI^-^/Ecad^-^/CK19^-^ DCC, respectively, occur *in vitro*, reflecting a developmental plasticity that may be controlled by a cell-autonomous process.

**Figure 4:**
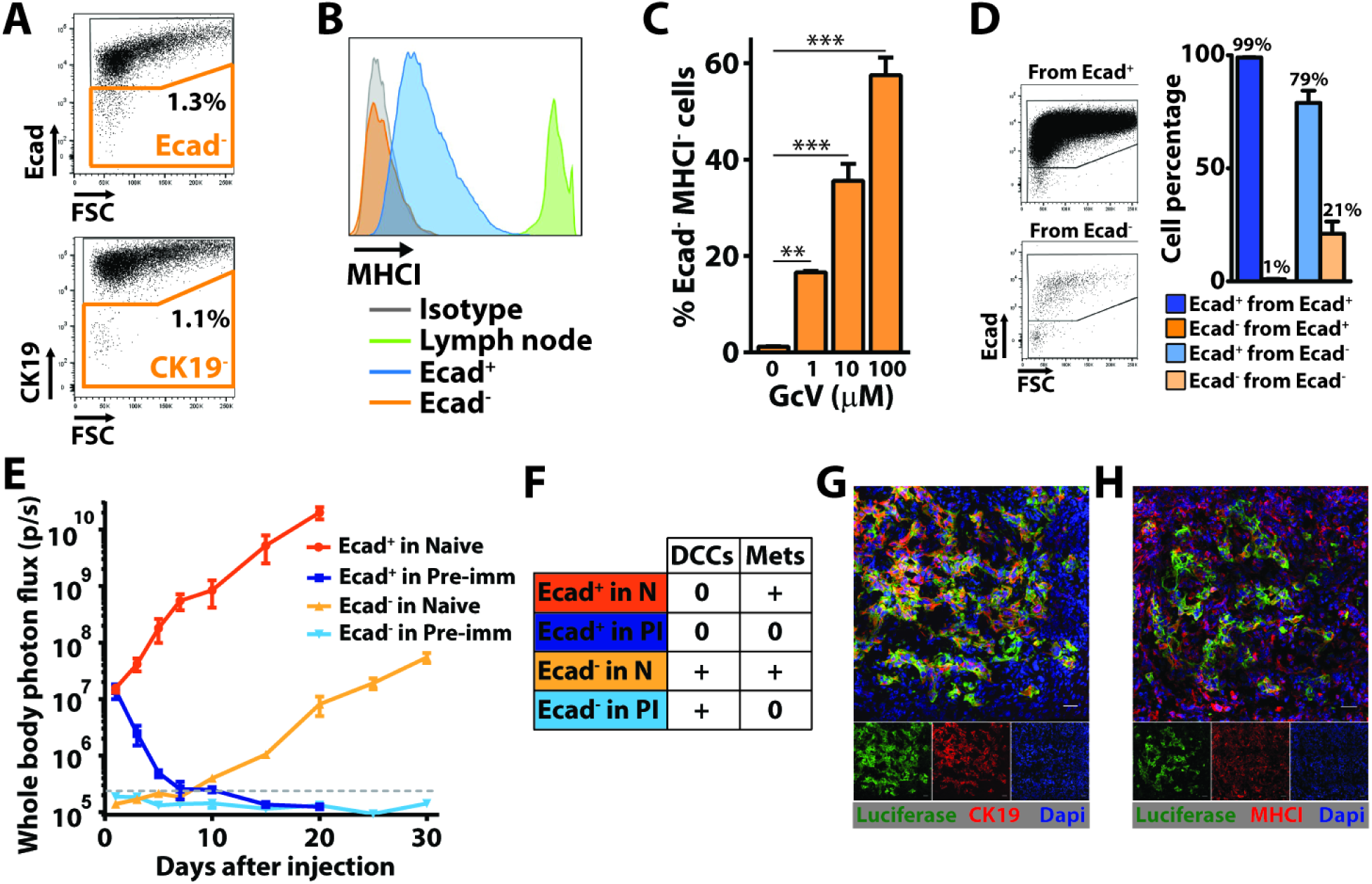
A subpopulation of PDA cells *in vitro* having the phenotype of DCCs. (A) Flow cytometry analysis of mM1DTLB PDA cells that have been stained with anti-CK19 or anti-Ecad. Results are representative of five independent experiments. (B) Flow cytometry measurement of anti-MHCI staining of Ecad^+^ and Ecad^-^ mM1DTLB PDA cells, and of lymph node cells as a comparator. Results are representative of five independent experiments. (C) mM1DTLB PDA cells were treated *in vitro* for 48h with increasing doses of GcV to kill proliferating cells, and the proportion of viable cells that was Ecad^-^/MHCI^-^ was measured by flow cytometry. Results are representative of two independent experiments. **=p<0.01, ***=p<0.001. (D) FACS analysis of purified Ecad^+^ and Ecad^-^ mM1DTLB PDA cells, respectively, that have been cultured for three days. Dot plots (left panel) and histograms (right panel) are representative of three independent experiments. (E) Growth of hepatic metastases after intra-splenic injection of 10^6^ Ecad^+^ or 10^4^ Ecad^-^ mM1DTLB PDA cells into naïve and pre-immunized mice was assessed by whole body bioluminescence imaging. n=5 mice per group. Dashed gray line represents the luminescence background in tumor-free mice. (F) Table summarizing the occurrence of DCCs and/or metastases in each group of mice. (G and H) IF of sections from a liver of a naïve mouse that had received an intra-splenic injection of Ecad^-^ mM1DTLB cells. Anti-luciferase (green) identifies cancer cells. Photomicrographs are representative of five mice. Scale bar=25μm.

### ER stress generates latent, immune-resistant DCCs

To identify the cell-autonomous “switch” regulating the developmental state of the metastases, we preformed single-cell RNA sequencing (scRNAseq) of *in vitro* sorted Ecad^+^ and Ecad^-^ cells. The most upregulated pathway in Ecad^-^ cells relative to Ecad^+^ cells is “Response to ER Stress” **(Fig. 5A and S9C)**, and the most downregulated pathway is “Cell Division” **(Fig. 5B and S9D)**. Network analysis of other upregulated pathways that distinguish the Ecad^-^ and Ecad^+^ PDA populations, such as “Autophagy”, shows that they are linked to “Response to ER Stress”. Similarly, network analysis of other downregulated pathways demonstrates linkage to “Cell Division” **(Fig. 5B)**. These two populations of the mM1DTLB PDA cells are also distinct by Principal Component Analysis (PCA), in which Ecad^-^ cells appear to be more heterogeneous than Ecad^+^ cells **(Fig. S9A)**. Indeed, analysis at the single cell level identifies four sub-populations of Ecad^-^ cells, in three of which the dominant upregulated pathway is related to “ER stress”, and the major downregulated pathway is “Cell Division” **(Fig. S10)**. An EMT signature was not present among the 1639 genes that were differentially expressed between Ecad^+^ and Ecad^-^ cells **(Fig. S9B)**, confirming the immunofluorescent analysis of Ecad^-^ cell **(Fig. S5E)**. Also, neither Ecad^+^ nor Ecad^-^ cells express three of the four major NKG2D ligands, Ulbp1, H60b and H60c, and strongly express the inhibitory ligand, Qa1, providing an explanation for the absence of a role for NK cells in controlling outgrowth of macro-metastases **(Fig. S6C)** (*26*). Genes that are involved in the processing of MHCI were not differentially expressed, which is consistent with reports that the ER stress response suppresses the expression of MHCI by a post-transcriptional mechanism (*27*, *28*). Finally, support for the conclusion that the ER stress response alters the expression of MHCI and Ecad is the finding that treatment of mM1DTLB PDA cells with tunicamycin increased the proportion of cells that were MHCI^-^and Ecad^-^ **(Fig. S11)**.

**Figure 5:**
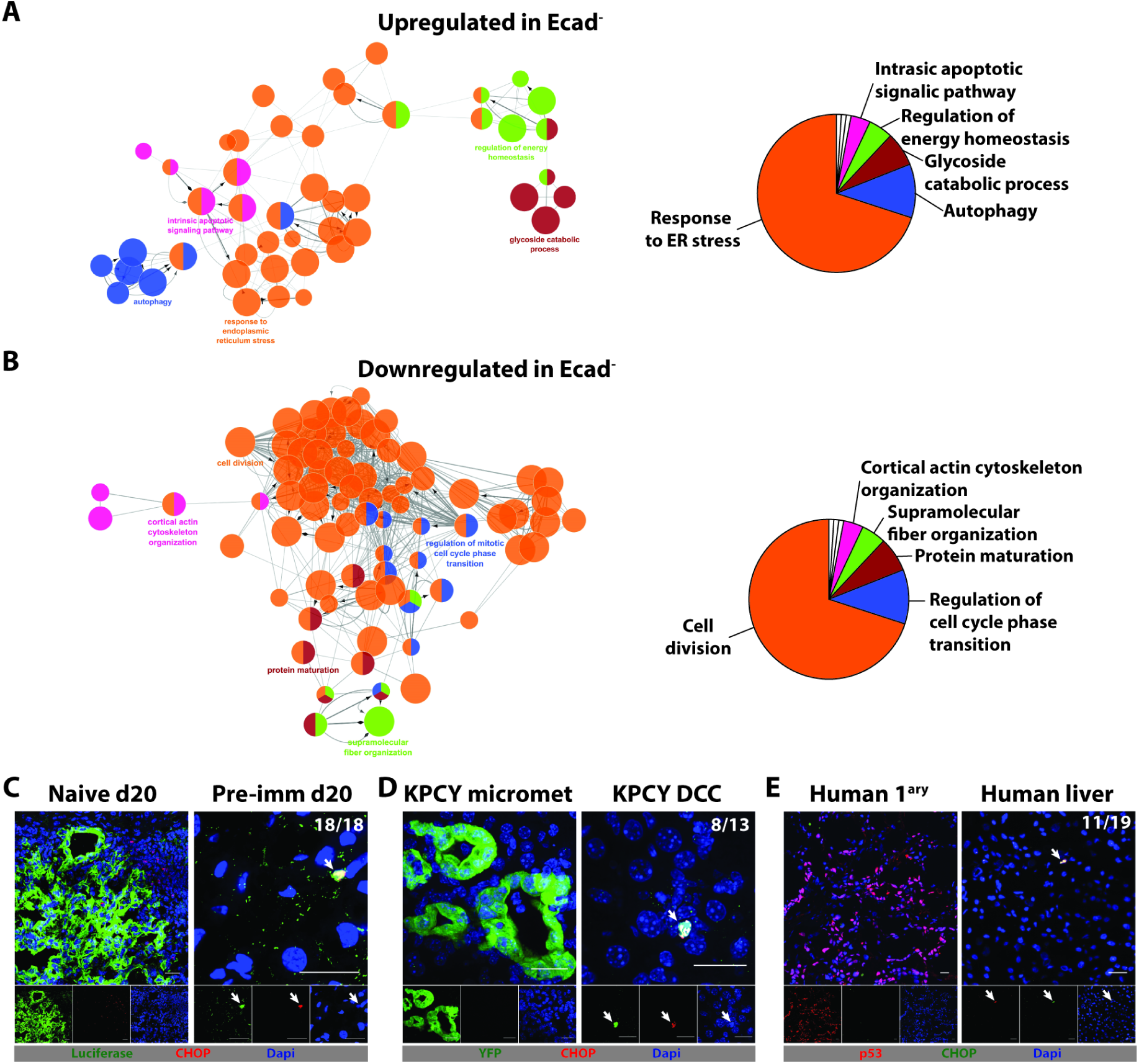
The ER stress response in PDA cells having the phenotype of DCCs. 104 Ecad^+^ and 98 Ecad^-^ mM1DTLB PDA cells were subjected to single cell RNA-Seq. (A) Network analysis, following pathway enrichment analysis, showing ontology relationships between the pathways (left panel) and their relative representation depicted as a pie chart (right panel) for (A) Upregulated pathways and (B) Downregulated pathways in the Ecad^-^ cells relative to Ecad+ cells. Pathways are significant with an adjusted p<0.01 after Benjamini-Hochberg false discovery rate. (C) IF of sections from a liver from a naive mouse and a pre-immunized mouse that have been stained with anti-luciferase (green) to identify PDA cells and anti-CHOP (red) to identify cells exhibiting an ER stress response. (D) IF of sections from a liver of a KPCY mice stained with anti-YFP (green) to identify PDA cells and anti-CHOP (red). (E) IF of sections from the primary tumor and liver of a patient with PDA that have been stained with anti-p53 to identify PDA cells (red) and anti-CHOP (green). Photomicrographs are representative of five patients, who had no detectable liver metastases and p53 loss-of-heterozygosity. Frequency of events within the DCC population is indicated in the top right corner of the photomicrographs. White arrows designate DCCs. Scale bar = 25μm.

The most differentially expressed gene was Ddit3/CHOP, the mRNA level of which was 18-fold higher in Ecad^-^ cells than in Ecad^+^ cells. CHOP is a transcription factor that is induced by the unfolded protein response (UPR), which is the basis of ER stress (*29*, *30*). We examined expression of CHOP protein by IF to determine whether the ER stress response also characterizes the Ecad^-^ DCCs in human and mouse primary and metastatic PDA. Anti-CHOP staining was demonstrated in hepatic DCCs in pre-immunized mice **(Fig. 5C)** and in KPCY mice **(Fig. 5D)**, but not in PDA cells of macro-metastases of naïve mice or of micro-metastases of KPCY mice (**Fig. 5E**). Most importantly, DCCs in the livers of 3/5 patients with PDA also stained with anti-CHOP antibody **(Fig. 5E)**. Thus, DCCs in both mouse and human PDA metastases may exhibit an ER stress response.

The chemical chaperone, 4-phenylbutyrate (4-PBA), binds to solvent-exposed hydrophobic segments of unfolded or improperly folded proteins, thereby “protecting” them from aggregation and relieving ER stress (*31*). Overnight treatment of mM1DTLB PDA cells with 4-PBA decreased the proportion of cells lacking expression of Ecad **(Fig. 6A)**, and increased MHCI expression in both Ecad^+^ and Ecad^-^ cells **(Fig. 6B)**. We also assessed the role of ER stress on proliferative capability of the cells. After eliminating the proliferating mM1DTLB PDA cells by treatment with GcV, the residual quiescent cells were pulsed with EdU overnight in the presence or absence of 4-PBA. Relieving ER stress with 4-PBA increased the proportion of cells incorporating EdU by 10-fold **(Fig. 6C)**.

**Figure 6:**
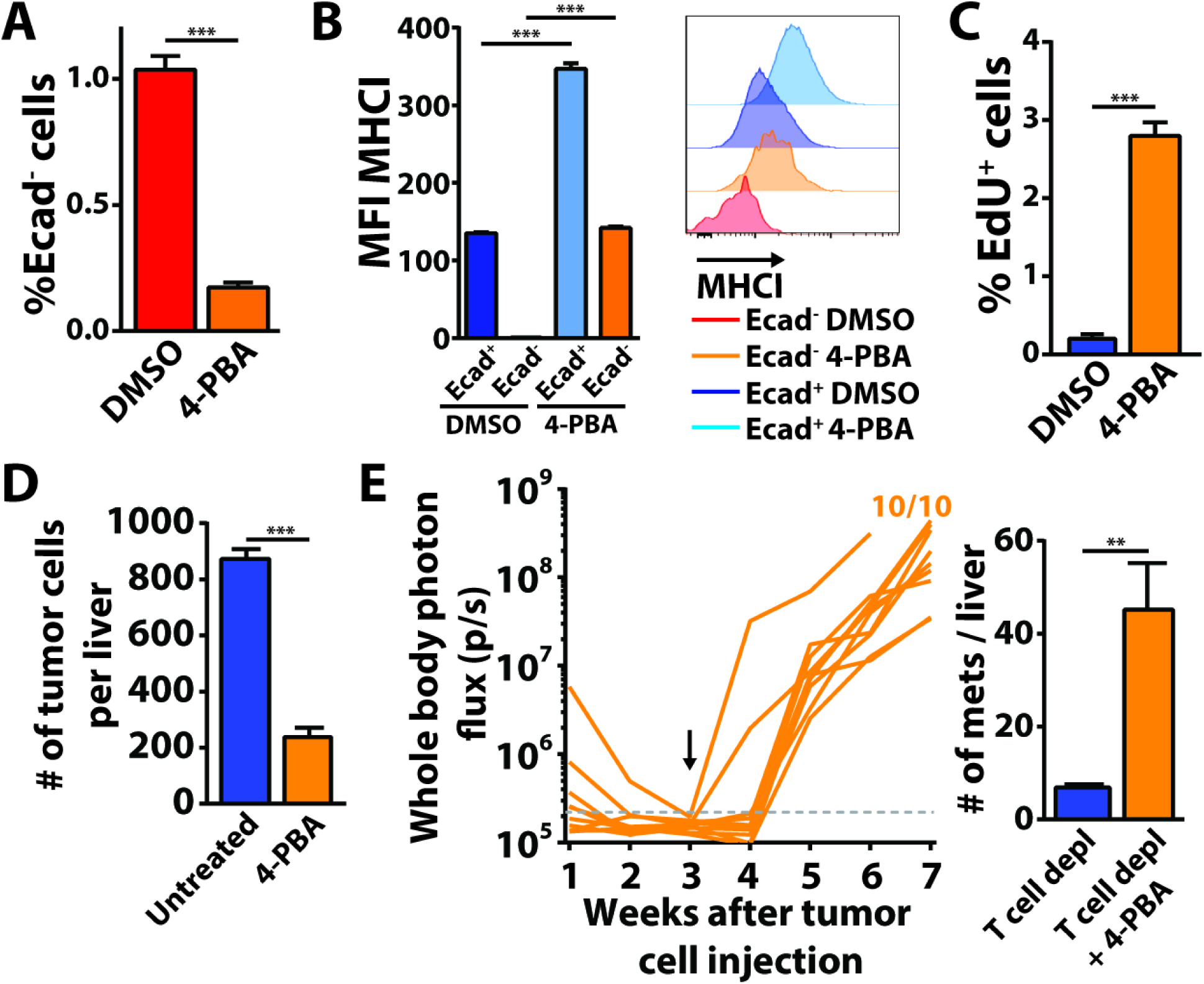
The effects on DCCs of relieving ER stress with a chemical chaperone. (A and B) mM1DTLB PDA cells were treated for 14 h with the chemical chaperone, 4-PBA, or DMSO. (A) The proportion of PDA cells that were Ecad^-^, (B) and the MFI of MHCI expression were measured by flow cytometry. Data are representative of three independent experiments. (C) mM1DTLB PDA cells were treated with 100μm GcV for two days and the remaining, non-cycling cells were pulsed overnight with EdU with or without 4-PBA. The proportion of EdU^+^ cells was assessed by flow cytometry. (D) Pre-immunized mice (n=5) that had received splenic injections of mM1DTLB PDA cells were treated with 4-PBA from the day of the injection. The number of hepatic DCCs was quantified after three weeks and compared to that of mice (n=5) not receiving 4-PBA. (E) Pre-immunized mice that had received splenic injections of mM1DTLB PDA cells were treated with 4-PBA beginning three weeks later. T cells were depleted by administering anti-CD4 and anti-CD8 antibodies. Growth of hepatic metastases was assessed by whole body bioluminescence imaging (left panel) and the number of metastases formed with or without 4-PBA treatment was counted by bioluminescence of the resected livers (right panel). n=10 mice. ** = p<0.01, *** = p<0.001.

We extended the analysis of the role of the ER stress response in DCCs *in vivo.* Pre-immunized mice were continuously treated with 4-PBA beginning on the day of the mM1DTLB cell injection, and livers were assessed for seeding by DCCs three weeks later. 4-PBA decreased number of DCCs by 4-fold **(Fig. 6D)**. The residual DCCs no longer demonstrated staining with anti-CHOP, and showed re-expression of MHCI **(Fig. S12A, B)**. The decrease in hepatic DCCs is consistent with the possibility that some DCCs in which ER stress had been relieved by 4-PBA had initiated a metastatic growth program and became targets for cytotoxic CD8^+^ T cells. To confirm this interpretation, we began 4-PBA treatment of pre-immunized mice three weeks after the intra-splenic injection of mM1DTLB cells together with T cell depletion. Within two weeks of 4-PBA treatment, all mice had developed MHCI^+^ macro-metastases **(Fig. 6E; Fig. S12C)**, and the number of macro-metastatic lesions in the livers of 4-PBA-treated, T cell-depleted mice was 6-fold higher than in the livers of mice subjected only to depletion of T cells **(Fig. 6E)**. These results are consistent with the ER stress response having a non-redundant, cell-autonomous role in the maintenance of quiescent, immune-resistant DCCs.

## Discussion

This clinical observation that PDA metastases develop in the majority of patients following the surgical removal of their primary tumors, despite no evidence of macro-metastases at the time of surgery, indicates that these patients harbored latent metastatic lesions. The nature of these latent metastases was suggested by our finding of single DCCs in the livers of patients and KPCY mice with PDA (**Fig. 1A-F**) that have a distinctive phenotype of absent CK19, E-cad, and MHCI. The absence of two typical markers of the epithelial ductal adenocarcinoma cells, without the occurrence of characteristic markers of EMT (**Fig. S2**), indicates that these DCCs are distinct from the PDA cells that comprise growing macro-metastases (*32*). Moreover, the absence of MHCI implies an unusual relationship of DCCs to the adaptive immune system (*33*). Although only descriptive, these findings provided the rationale for developing a mouse model that replicates DCCs with this distinctive phenotype and allows mechanistic studies defining both the cell-autonomous response responsible for the phenotype, and the role of the immune system.

We hypothesized that, at least in some patients with PDA, metastases occur in the context of a cancer-specific adaptive immune response (*19*-*22*). The absence of MHCI on DCCs in human and mouse PDA, and the absent or infrequent occurrence of hepatic macro-metastases in which cancer cells express epithelial markers and MHCI, raised the possibility that immunity prevents the outgrowth of macro-metastases while ignoring MHCI^-^ DCCs. This prediction was based also on the observation that DCCs in human and mouse PDA, in contrast to cancer cells in macro-metastases, were quiescent and infrequently Ki67^+^ (**Fig. 1**). The finding that pre-existing immunity prevented the occurrence of hepatic macro-metastases while permitting the seeding of non-replicating, MHCI^-^ DCCs throughout the liver verified this prediction, and provided a potential explanation of how quiescent metastases can persist in the presence of an adaptive immune response that is capable of suppressing the growth of macro-metastases.

This selective effect of adaptive immunity on macro-metastases required that the cell-autonomous mechanism that is responsible for the phenotype of DCCs invariably links the expression of MHCI to a capacity for cellular replication. This linkage was not only demonstrated in additional cell lines from primary and metastatic PDA tumors from KPC mice (**Fig. S3, S4**), but also was supported by the finding that ER stress, which inhibits MHCI expression (*27*, *28*), and cell division (*34*), were the major up-regulated and down-regulated transcriptional signatures that distinguish Ecad^-^ cells from Ecad^+^ PDA cells. Thus, MHCI expression is “off” in quiescent cells and “on” in replicating cells. In addition, if the markedly increased expression of CHOP in Ecad^-^ PDA cells, which is a transcription factor that is induced by the UPR (*29*, *30*), is taken as an indicator of the ER stress response, then the finding that in human and mouse PDA, hepatic DCCs are MHCI^-^, Ki67^-^ and CHOP^+^ **(Figs. 1 and 5)** links quiescence and immune concealment to the ER stress response *in vivo*.

A latent capacity of hepatic DCCs for reversion to growing cancer cells was revealed by the outgrowth of macro-metastases following the depletion of T cells (**Fig. 3**). This finding also implied that T cells eliminated those DCCs stochastically reverting to a replicating, MHCI^+^, epithelial phenotype. T cell killing of reverting DCCs must be efficient and occur before an immune suppressive microenvironment is established since no macro-metastases were observed during a 12 month period of observation of T cell-replete mice with hepatic DCCs (**Fig. 3**). The causal link between reversal of the DCC to a replicating, MHCI^+^, epithelial phenotype was established by the use of the chemical chaperone, 4-PBA. Relief of ER stress by 4-PBA enhanced the expression and E-cad and MHCI by DCC-like cells while promoting their proliferation *in vitro* (**Fig. 6**), and, most importantly, caused the outgrowth of macro-metastases in T cell-depleted mice bearing hepatic DCCs. This observation, coupled with the capacity of 4-PBA to decrease hepatic DCCs in T cell-replete mice, supports the essential role of the ER stress response in maintaining the DCC phenotype.

The implications for therapy to prevent the occurrence of metastatic disease in patients following the surgical removal of their primary PDAs may be two-fold. First, outgrowth of latent DCCs in the mouse model requires suppression of T cell immunity (**Fig. 3**). Elevations of plasma cortisol following pancreatectomy in patients with PDA (*35*) are in the range that was found to be immune suppressive in cachectic mice with PDA (*36*). This stimulation of the hypothalamic-pituitary-adrenal axis may also occur with the caloric deprivation that commonly occurs in patients after this surgical procedure (*37*). Therefore, post-operative parenteral hyper-alimentation may be an effective means to decrease the occurrence of metastatic disease in patients following surgical removal of their primary PDAs. Second, and perhaps more speculative, the administration of a chemical chaperone, like 4-PBA, pre-operatively when tumor immunity is intact might be an approach to purge organs of latent DCCs, thereby decreasing likelihood of postoperative metastatic disease.

## Acknowledgments

We thank Dr. David Tuveson for providing mM1 and 1242 PDA cells as well as the KPCY liver tissues. **Funding:** This work was supported by a Distinguished Scholar Award to D.T.F. from the Lustgarten Foundation, an award from the Cedar Hill Foundation, and 5P30CA45508-29, NIH-NCI. A.P. was supported by the Philippe Foundation. **Competing interests:** The authors disclose no potential conflicts of interest.

## Supplementary materials

Materials and methods

Supplementary figures S1 to S12

Supplementary references

